# High levels of DNA replication initiation factors indicate ATRi sensitivity via excessive origin firing

**DOI:** 10.1101/2025.02.27.640046

**Authors:** A Lumeau, PL Pfuderer, S De Angelis, JA Scarth, MA Guscott, N Shaikh, FB Copley, H Gerdes, EL Alard, PR Cutillas, F Mardakheh, MA Boemo, JV Forment, SE McClelland

## Abstract

Inhibitors of ATR, a central kinase controlling DNA replication origin firing and cellular checkpoint activity, are currently in multiple clinical trials, yet mechanisms underpinning sensitivity and robust patient stratification biomarkers are lacking. We used functional genomics approaches to identify molecular mechanisms driving sensitivity to the ATR inhibitor (ATRi) ceralasertib. Replication stress-associated patterns of DNA copy number alterations identified a subset of sensitive breast cancer cell lines. In parallel, we performed proteomics, phosphoproteomics and gene expression analyses and discovered that sensitive cell lines had higher expression of DNA replication origin firing factors, and massively increased origin firing in response to ATRi. ATRi sensitivity was partly rescued upon co-treatment with XL-413, a CDC7 inhibitor that decreases origin firing. High expression of replication initiation factors correlated with ATRi sensitivity across multiple cancer types, and in acute myeloid leukemia patient samples. Together, this study reveals a novel contribution of lethal origin firing capacity in determining the sensitivity of cancer cells to ATR inhibition and demonstrates the predictive potential of mechanism-specific copy number alterations, providing key steps towards developing a multimodal clinically applicable biomarker for ATR inhibitors.

## Introduction

Replication stress (RS) is defined as the slowing or stalling of DNA replication forks and has been shown to be driven by oncogene activation across several tumor types^1,2^. RS is a known driver of chromosomal instability (CIN), characterized by continuous gains and losses of whole or parts of chromosomes during cell division, in multiple cancer types^3–5^. Cell cycle checkpoints, the RS response and DNA repair pathways are crucial for alleviating RS in normal cells, ensuring accuracy and fidelity during cell division to limit CIN. A common cancer therapeutic strategy is to capitalize on the high levels of RS, genomic instability and/or cell cycle checkpoint defects in cancer cells by inhibiting these protective pathways^6^. Accordingly, several RS response inhibitors are in clinical trials based on the principle that higher underlying rates of RS will sensitize tumors to a further loss of key RS responses or checkpoints^6,7^.

ATR (Ataxia telangiectasia and Rad3 related) is a protein kinase that maintains genomic integrity by regulating origin firing during normal S-phase, and by acting at cell cycle checkpoints to limit entry into mitosis in the presence of RS or DNA damage^8^. ATR inhibition therefore has two major consequences for the cell. Firstly, unscheduled origin firing and abrogation of the intra-S-phase checkpoint induces widespread RS^9,10^. Secondly, when the G2/M checkpoint is impeded, cells with damaged genomes can continue into mitosis, potentially causing unrestrained genomic instability^11^.

ATR kinase inhibitors are an emerging new class of anti-cancer treatment being developed to exploit elevated RS, DNA damage response deficiencies and/or to promote anti-tumor immunity in cancer^11,12^. Multiple ATR inhibitors are in clinical trials as monotherapy or in combination with a diverse range of partner agents and across multiple cancer types (reflecting the complex biology of ATR), although none are yet approved. To date, clinical monotherapy efficacy has been modest and can be associated with dose-limiting toxicities (particularly hematological). Encouraging clinical activities have been recently reported for ATRi in combination with other therapies; for example Phase 2 clinical studies combining ceralasertib with the Poly (ADP-ribose) Polymerase (PARP) inhibitor olaparib in PARPi-resistant ovarian cancer patients^13^ or with the anti-PD-L1 immune checkpoint inhibitor durvalumab in immunotherapy-resistant non-small cell lung cancer patients^14^, but no standout patient selection biomarker has been identified^11,15^. It is therefore crucial to improve understanding of the mechanism of action of ATRi, to identify biomarkers to predict patient response, as well as revealing more effective therapy combinations.

The identification of biomarkers has been largely focused on transcriptomic signatures. Genetic and genomic signatures are an alternative approach and have been employed in predicting sensitivity to PARP inhibitors in patients who lack homologous recombination repair ability (e.g. due to BRCA1/2 mutations^16^). Our previous work determined that patterns of genomic alterations can reflect the underlying mechanisms of genomic instability, and defined copy number alteration features specific to RS induction^17^. Moreover, pan-cancer efforts to characterize patterns of genomic alteration from tumors to predict underlying CIN mechanisms have also demonstrated that RS is likely a key driver of CIN in cancer that could indicate sensitivity to RS response inhibitors^18,19^.

In this study, we aimed to understand the mechanisms contributing to ATRi sensitivity, and to explore multimodal biomarkers that could be used in future clinical trials involving ATRi. We confirmed a general link between increased RS and ATRi sensitivity in non-transformed cell lines and demonstrated that sensitive breast cancer cell lines exhibit lower replication fork speeds. Moreover, we could predict a subset of sensitive cell lines by RS-specific copy number alteration patterns extracted from our previous study^17^. However, all cell lines experienced similar increases in RS and CIN upon ATR inhibition, challenging the assumption that ATRi causes intolerable rates of RS and/or CIN, and prompting us to explore the precise molecular mechanisms responsible for cell death in sensitive cell lines. We performed an unbiased approach, using proteomics and gene expression analyses and discovered that sensitive cell lines expressed higher levels of replication initiation factors. Functionally, sensitive cell lines exhibited hyper-increases in origin firing upon ATRi treatment, and ATRi sensitivity was reproducibly reduced by lowering origin firing capacity. ATRi-sensitive cell lines from multiple cancer types and patient-derived AML primary cell lines exhibited significantly enriched expression of DNA replication initiation factors, providing a potential new indicator of ATRi sensitivity.

## Results

### Replication stress is associated with sensitivity to ATRi

First, to validate the association between RS and ATRi sensitivity, we used two h-TERT immortalized, non-transformed diploid cell lines; RPE1 (retinal pigment epithelial cells) and FNE1 (fallopian tube epithelial cells)^20^. RPE1 cells treated with low doses of the DNA polymerase poison aphidicolin that nonetheless permit cell cycle progression^17^ showed elevated DNA damage in replicating cells (as measured by phosphorylation of histone variant H2AX on Ser-139, also known as γH2AX; **Extended Figure 1a**), increased micronuclei resulting from CIN (**Extended Figure 1b**) and were sensitized to ATRi treatment (**Figure 1a, Extended Figure 1c**). FNE1 cells treated with aphidicolin, or after induction of *CCNE1* (Cyclin E1) overexpression (**Extended Figure 1d**), previously implicated as an oncogenic driver of RS^21–23^, were also sensitized to ATRi (**Figure 1b**), thus confirming a relationship between RS and ATRi sensitivity.

**Figure 1.**
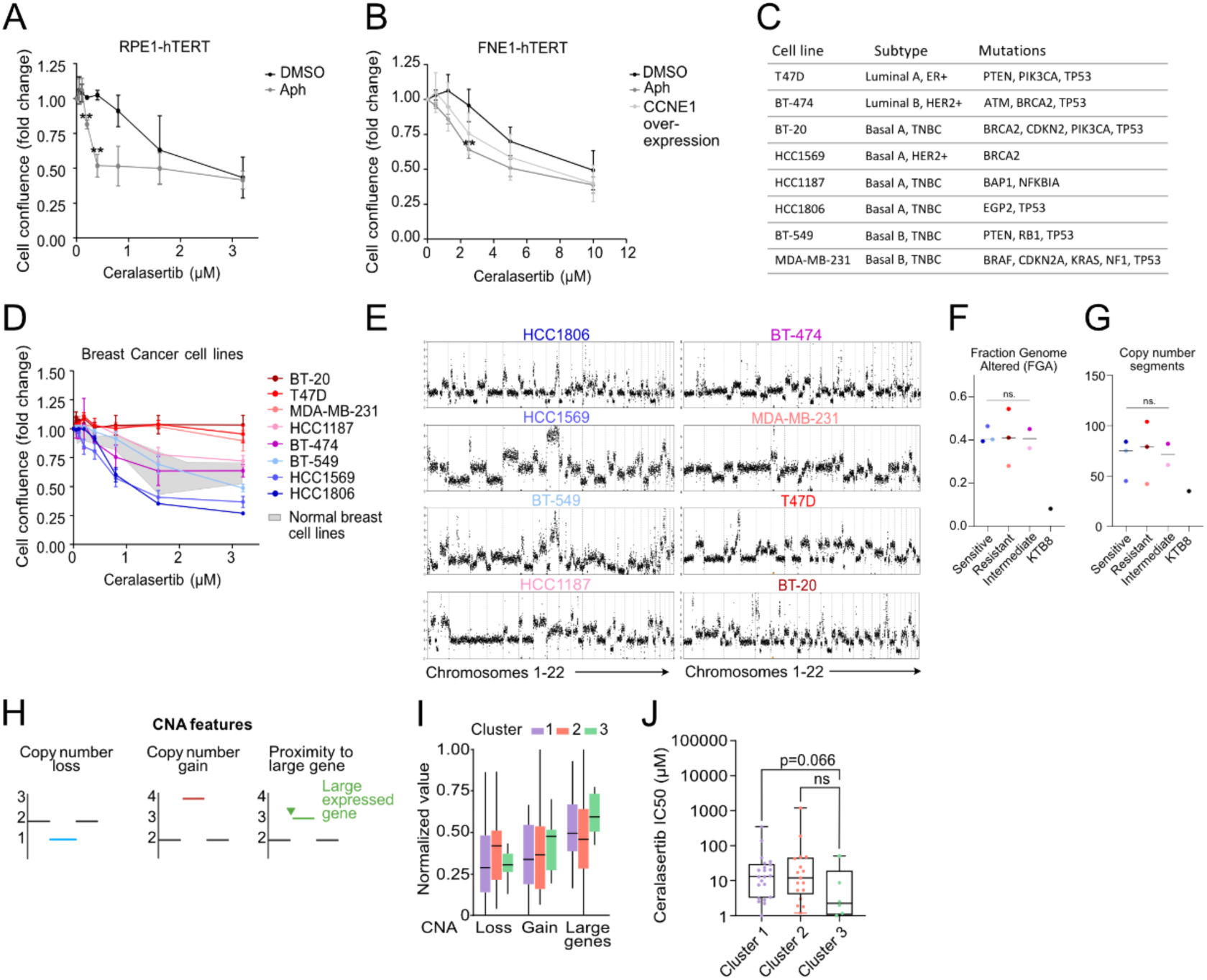
General replication stress is associated with sensitivity to ATRi. (**A,B**) Cell confluence after 72h at the doses of ceralasertib (ATRi) indicated in RPE1 hTERT co-treated with DMSO or Aphidicolin 0.4µM (A) or FNE1 hTERT cells overexpressing CCNE1 by tetracycline induction 48h pre-ATRi treatment (B). Multiple paired t-test analysis. (**C**) Table indicating the genetic and subtype details of a panel of breast cancer cell lines. (**D**) Cell confluence after 72 h at the doses of ceralasertib (ATRi) indicated in the breast cancer cell lines. For A), B) and D) fold confluence of ATRi treated samples was measured relative to DMSO at 72h. Results are representative of three independent experiments for every cell line. HCC1806, HCC1569 and BT-549 are classified herein as the most sensitive, HCC1187 and BT-474 as intermediate and T47D, MDA-MB-231 and BT-20 most resistant. (**E**) Representation of the genomic copy number plots obtained from long read nanopore sequencing. The x-axis represents the 22 chromosomes and the y-axis the copy number. (**F,G**) Quantification of the fraction of genome altered (F) and the number of copy number segments (G) of the sensitive and resistant cell lines. Unpaired t-test analysis. (**H**) Schematic representation of the replication stress-associated CNA features approach. (**I**) Quantification of the normalized CNA value for three different CNA features: loss, gain, and proximity to large, expressed genes, into three different clusters. (**J**) Quantification of the IC50 in µM across the copy number features-defined clusters, using IC50 values from GDSC2 datasets. Unpaired t-test analysis. Ns, not significant; *, P < 0.05; **, P < 0.01.

Since ATR inhibitors are currently in clinical trials in many cancer types, including breast cancer, we chose to test the effects of ATRi on a panel of breast cancer cell lines. There is no known link between breast cancer subtype and ATRi sensitivity, so our panel included CCLE (Cancer Cell Line Encyclopedia) cell lines from luminal and basal breast cancer subtypes with different genetic alterations (**Figure 1c)**. The panel exhibited a range of ATRi sensitivities (**Figure 1d, Extended Figure 1c,e**) facilitating discovery of differences between sensitive and resistant lines. As a tissue-specific comparative control we also assessed the ATRi sensitivity of a panel of three non-transformed breast cell lines (KTB6, KTB8, and KTB34)^24^ (**Extended Figure 1g-h**). We classified three breast cancer cell lines as more resistant (BT-20, T47D, MDA-MB-231) and three as more sensitive (HCC1569, HCC1806, BT-549) to ATRi than the non-transformed KTB panel, with the remaining two (HCC1187, BT-474) showing an intermediate sensitivity phenotype (**Figure 1d**). Previous work has reported that ATRi sensitivity is associated with *CCNE1* (cyclin E) amplification in different cellular models^23,25^. However, using publicly available transcriptomic data and IC50 to ceralasertib of breast cancer cell lines, we found no correlation between ATRi sensitivity and expression of *CCNE1*, *CCNE2* or *MYC* (all potential oncogenic drivers of RS) (**Extended Figure 1f**).

We have previously demonstrated that chromosomal instability resulting from RS generates distinctive patterns of DNA copy number alterations (CNAs)^17^. We therefore hypothesized that an examination of the CNA landscape of a panel of breast cancer cell lines might reveal ATRi sensitivity-specific genomic features derived from inherent RS. Using DNA CNA analysis from Oxford Nanopore long-read whole genome sequencing, we observed that all lines exhibited chaotic genomes with gains and losses of whole and partial chromosomes (**Figure 1e**). Neither the fraction of genome altered, nor the number of DNA CNAs differed between sensitive and resistant lines (**Figure 1f,g**). We therefore analyzed a larger panel of breast cancer cell lines for which copy number profiles could be determined from publicly available exome sequencing data^26^, and clustered cell lines based on the presence of various RS-associated copy number features defined in our earlier study^17^, namely the frequency at which DNA copy number breakpoints occurred proximal to large (>600 kb), expressed genes, and the total fractions of genomic losses and gains (**Figure 1h,i, Extended Figure 1i**). We then categorized each cell line as sensitive or resistant to ATRi based on data from the “Genomics of Drug Sensitivity in Cancer” database^27^. This approach revealed one cluster of samples (Cluster 3) that contained mainly sensitive cell lines (**Figure 1j**), apart from one cell line HCC1187, demonstrating the potential of leveraging experimentally defined DNA CNA patterns caused by RS to predict ATRi sensitivity.

### ATRi sensitivity is not due to an intolerable increase in RS-associated chromosomal instability

To complement the genomic signature approach above and identify additional sensitive cell lines, we set out to gain more mechanistic insight into ATRi sensitivity. Since RS is a driver of CIN^28,29^, either RS or the resulting increase in CIN may be responsible for ATRi sensitivity. For the following analyses, the three most sensitive cell lines (HCC1806, HCC1569 and BT-549) were compared to the three most resistant (MDA-MB-231, T47D and BT-20) (**Figure 1c**). We hypothesized that cell lines demonstrating the highest levels of RS (and thus the slowest replication fork rates^1,30^) would be the most sensitive to ATRi. To directly assess RS levels, we examined DNA replication dynamics using a novel long-read sequencing method, DNAscent, which detects replication fork speed and stall frequency using machine learning to call base analogues in recently replicated DNA^31^ (**Figure 2a**). In baseline conditions (before ATRi treatment), neither replication fork speeds nor fork stalling rates were significantly different between sensitive and resistant cell lines (**Figure 2b-e**). Furthermore, there was no significant difference in the level of S-phase γH2AX foci, a global mark of replication-associated DNA damage, between the sensitive or resistant cell lines (**Extended Figure 2a**). Since ATR inhibition impacts both origin firing rates and RS response signaling, the effect on replication dynamics may differ over time. Therefore, we performed DNAscent after 1, or 24 hours ATRi treatment to capture both the initial and longer-term impact of ATR inhibition on DNA replication kinetics. In response to ATRi, all cell lines showed a significant decrease in fork speed, an increase in fork stalling, or both, demonstrating increased RS regardless of sensitivity status (**Figure 2b,d**). Overall, therefore, canonical biological markers of RS could not robustly distinguish ATRi sensitive cell lines.

**Figure 2.**
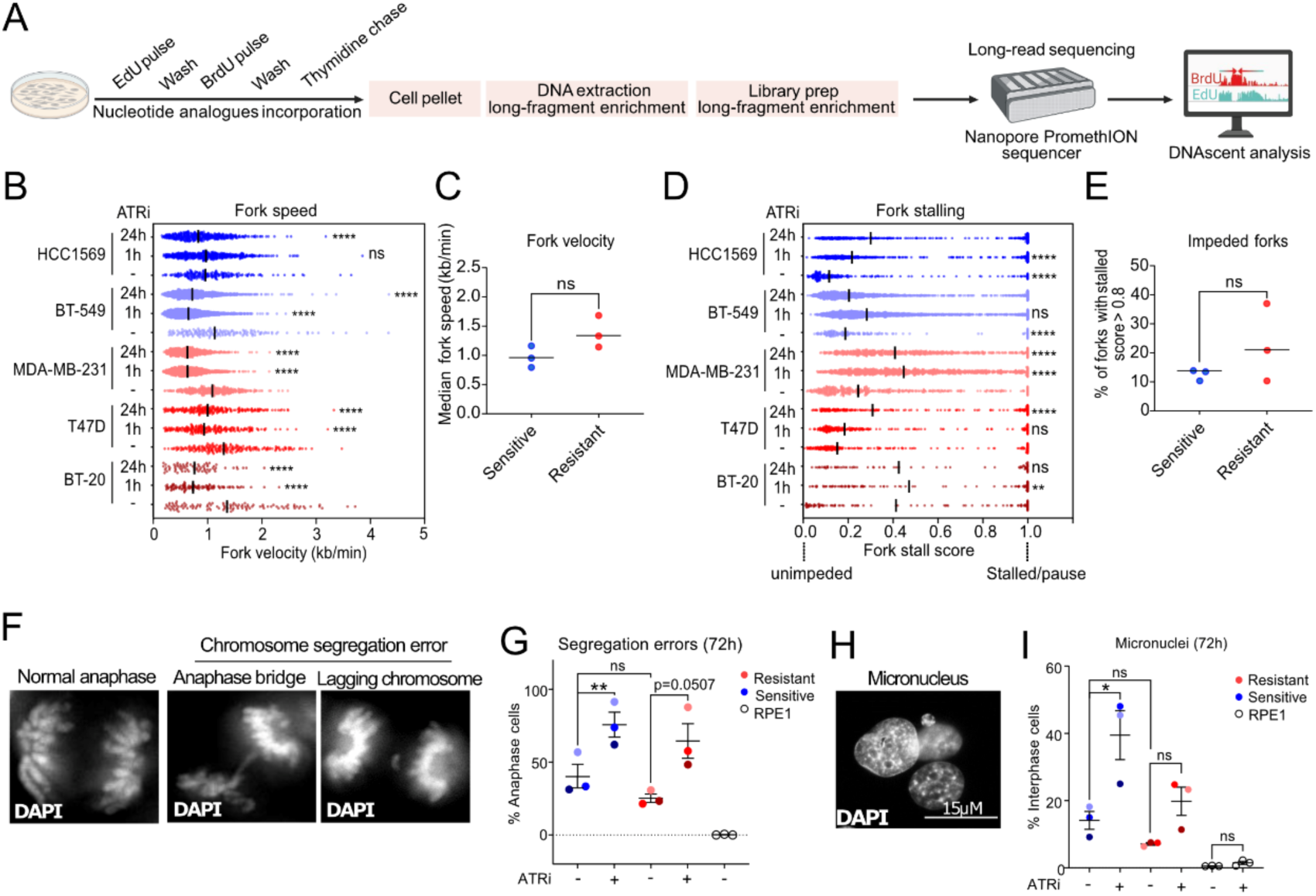
ATRi sensitivity is not due to an intolerable increase in replication stress-associated chromosomal instability. (**A**) Schematic representation of the method to generate replication fork dynamics via nucleotide analogue incorporation, nanopore long read sequencing and DNAscent algorithms. Quantification of the fork (**B**) velocity (kb per minute), (**C**) the median fork velocity (kb per min), (**D**) the fork stalling probability and (**E**) the proportion of impeded forks (with a stall score probability above 0.8) in sensitive and resistant cell lines treated either with DMSO or 1h or 24h of 0.5 µM ATRi. (B,D) Mann & Whitney analysis between each ATRi and DMSO per cell line. (C,E) Unpaired t-test analysis between three sensitive and three resistant cell lines. Note that for this experiment, only the DMSO condition was performed for HCC1806. (**F**) Representative microscopy images of a normal anaphase, an anaphase bridge and a lagging chromosome. Chromatin visualized with DAPI. (**G**) Quantification of the segregation error rate in cells treated with DMSO or ATRi 0.5 µM for 72 hours. (**H**) representative image of an interphase cell with a micronucleus. (**I**) Quantification of the micronuclei rate in cells treated with DMSO or ATRi 0.5 µM for 72 hours. Results represent the average of three sensitive cell lines compared to three resistant cell lines with at least two independent replicates per cell line. Paired t-test analysis. Ns, not significant; *, P < 0.05; **, P < 0.01; ****, P < 0.0001.

We next determined whether rates of RS-induced CIN were different between sensitive and resistant lines, by scoring the rates of chromosome mis-segregation and micronuclei before and after treatment with ATRi. Baseline rates of CIN were not significantly different between the sensitive and resistant lines (**Figure 2f-i, Extended Figure 2d**). Furthermore, although all cell lines displayed increased CIN after ATRi treatment, there were no clear differences in the rates nor type of CIN induced by 72h ATRi treatment between sensitive and resistant cell lines (**Figure 2f-i, Extended Figure 2b-d**). The universal increase in CIN following ATRi treatment also excludes differences in drug uptake or efflux as a cause of differential ATRi sensitivity. Therefore, neither high baseline rates of CIN nor intolerable levels of ATRi-induced RS or instability could explain the differential ATRi sensitivity of this panel of breast cancer cell lines.

We and others have previously shown that supplementing cells with exogenous nucleosides can reduce RS and CIN^28,29,32^. In line with the absence of strong associations between RS or CIN levels and ATRi sensitivity in these cell lines, supplementing cells with nucleosides only very mildly rescued the sensitivity in HCC1806 (**Extended Figure 2e**), suggesting other underlying mechanisms of sensitivity to ATRi.

### Increased origin firing capacity renders sensitive cell lines more vulnerable to cell death following ATR inhibition

To search in an unbiased manner for the specific mechanisms causing differential ATRi sensitivity, we performed total and phosphoproteomic analyses using tandem mass tagging (TMT) mass spectrometry to allow quantitative comparisons between sensitive and resistant cell lines (**Figure 3a**).

**Figure 3.**
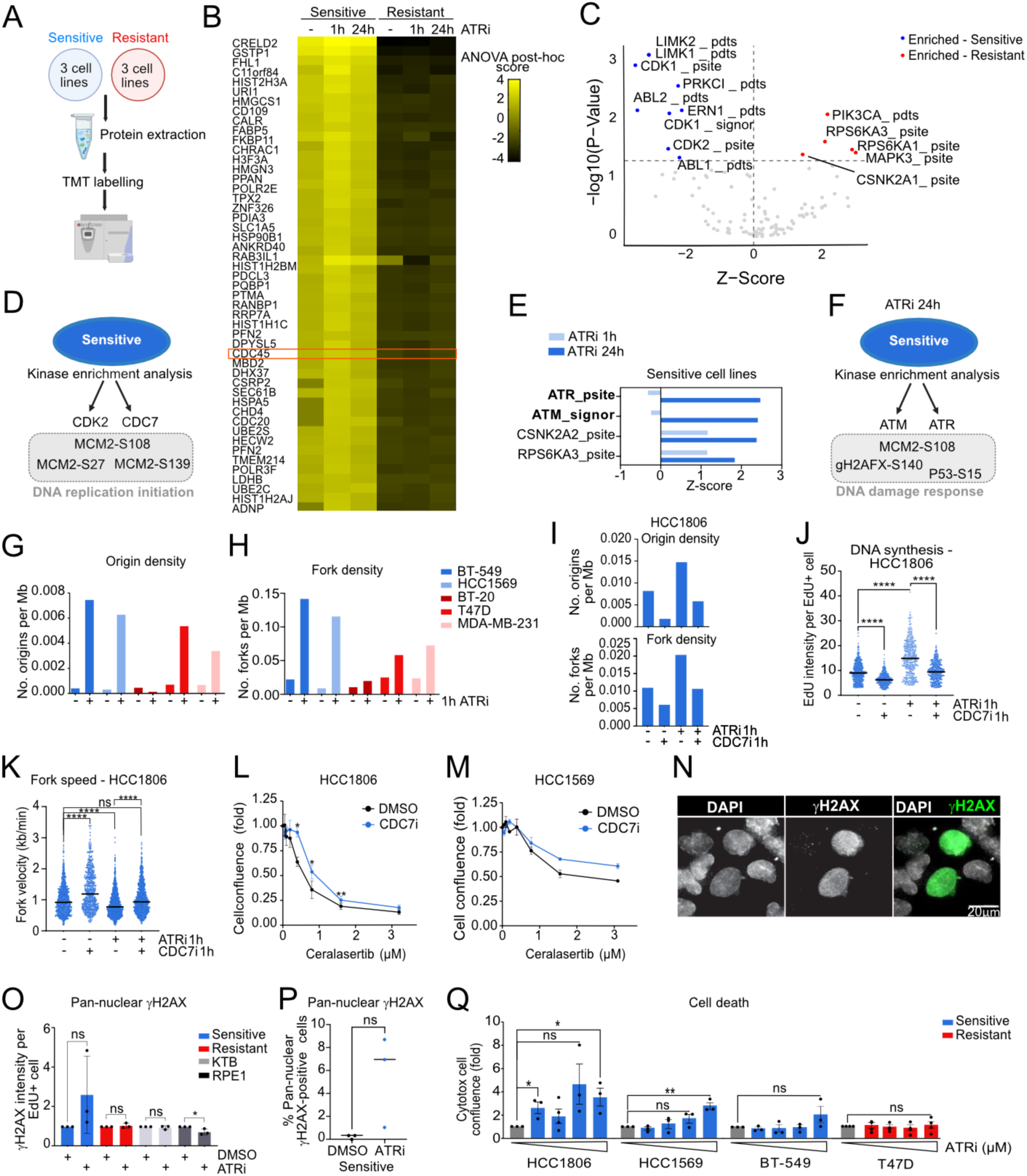
Increased origin firing capacity renders cancer cell lines more vulnerable to ATR inhibition. (**A**) Schematic representation of the mass spectrometry method to obtain total proteome and phosphoproteomes of breast cancer cell lines. (**B**) Heatmap plot showing the top hit significantly enriched proteins in sensitive compared to resistant cell lines. The orange box shows CDC45 protein. Significantly changing proteins were identified using a one-way ANOVA, with post-hoc score calculation (FDR < 0.05). (**C**) Volcano plot representation of the kinase activities most enriched in sensitive and resistant breast cancer cell lines, obtained by kinase enrichment analysis of phosphoproteomics data measured by mass spectrometry. (**D**) Schematic representation of the enrichment of kinases CDK2 and CDC7 in sensitive cell lines at baseline conditions, exemplified by an increase of phosphorylation of MCM2 at sites S108, S27 and S139, involved in origin firing. (**E**) Quantification of kinase activities enriched in response to 24 hours of 0.5 µM ATRi in sensitive cell lines compared to resistant, Z-scores higher than 1.5. (**F**) Schematic representation of the enrichment of kinases ATM and ATR via activation of substrates MCM2 S108, γH2AX-S140 and p53 S15 involved suggesting a strong activation of the DNA damage response pathway. (**G,H**) Quantification of the replication origins (**G**) and fork density per megabase (**H**) in breast cancer cell lines treated with DMSO or 0.5 µM ATRi for 1 hour. Data obtained from R9 long-read DNA sequencing. (**I**) Quantification of the replication origins and fork density per megabase in sensitive HCC1806 cells, either with DMSO, 0.5 µM ATRi or 1 µM CDC7i or combination for 1 hour. Data obtained from R10 long-read DNA sequencing. (**J**) Quantification of the mean EdU intensity in replicating cells (EdU+) in HCC1806 (sensitive cell line) cells treated with DMSO, 1 µM CDC7 inhibitor (XL-413), 0.5 µM ATRi or the combination for 1 hour. EdU intensity is used as a metric of activation of origin firing. Results presented are representative of two independent replicates with 250 to 500 EdU positive cells quantified per condition. Paired t-test analysis. (**K**) Quantification of the fork velocity (kb per minute) in sensitive HCC1806 cells treated either with DMSO, 0.5 µM ATRi or 1 µM CDC7i or combination for 1 hour. Data obtained from R10.4.1 long-read DNA sequencing (see methods). Mann & Whitney analysis between each treatment condition and DMSO. (**L,M**) Quantification of the dose response to ATRi in combination with DMSO or 0.25 µM of CDC7i in HCC1806 (L) and HCC1569 (M) sensitive breast cancer cell lines. Multiple paired t-test analysis. (**N**) Microscope imaging of γH2AX pan-nuclear staining in the cell population. (**O**) Quantification of γH2AX intensity in replicating cells (EdU+) in response to DMSO or 0.5 µM for 24 hours in breast cancer cell lines, KTB cells and RPE cells as indicated. Paired t-test analysis. (**P**) Quantification of the percentage of cells with pan-nuclear γH2AX staining in the three sensitive cell lines treated with DMSO or 0.5 µM ATRi for 24 hours. Paired t-test analysis. (**Q**) Quantification of the green Cytotox confluence, live death marker, as a fold ratio to DMSO in HCC1806, HCC159, BT-549 and T47D treated with DMSO or 72h ATRi (0.4 µM, 0.8 µM, 1.6 µM, 3.2 µM). Paired t-test analysis. Ns, not significant; *, P < 0.05; **, P < 0.01; ****, P < 0.0001.

For this, we treated the three most resistant or sensitive breast cancer cell lines with DMSO or 0.5 µM of ATRi for 1 h or 24 h before TMT, detecting 5773 phosphosites across 5000 proteins. Post-hoc scores were measured using an ANOVA analysis across all conditions. Under baseline (untreated) conditions, CDC45, a key regulator of replication origin firing^10,33^, was among the top 50 enriched proteins in sensitive compared to resistant lines (**Figure 3b)**. We therefore searched for additional proteins in the origin firing pathway using the 37 gene ‘‘DNA replication initiation” GO term. Of the 20 members of this GO term detected in our dataset, three additional origin firing proteins (ORC1, KAT7, WRNIP1) were found to be enriched in sensitive versus resistant cell lines (**Extended Figure 3a**). We also confirmed the increase of *CDC45* expression in sensitive cell lines at the mRNA level using publicly available data (**Extended Figure 3b**). Turning next to the phosphoproteome, we searched for differences in the cellular signaling consequences of ATRi treatment between the sensitive and resistant lines. Unfortunately, we could not directly identify several key phosphorylation sites of interest, including the CHK1 serine 345 phosphorylation site, direct substrate of ATR^34^. We therefore used a Kinase Enrichment Analysis Method (KSEA), which allows the inference of activated kinases from the presence of downstream target phosphorylation events^35^. Using this analysis, we noted that CDK2 and CDC7 signaling pathways were enriched under baseline conditions in sensitive cell lines compared to resistant cell lines (**Figure 3c, Extended Figure 3c**). CDK2 and CDC7 are involved in origin firing, including via MCM2 phosphorylation sites that were found enriched in sensitive cells (**Figure 3d**). In addition, only sensitive cell lines exhibited evidence of increased DNA damage signaling upon 24h of ATRi treatment, with increased ATR and ATM pathway activation, exemplified by the specific phosphorylation of MCM2, γH2AX and p53 (**Figure 3e-f, Extended Figure 3d**). Note that using this analysis approach, ATR signaling pathways are detected as active despite the presence of ATRi due to the overlap in substrates between the ATR and ATM signaling pathways^36^. Overall, these data suggested that sensitive cells were experiencing higher levels of origin firing under untreated conditions, as well as higher levels of DNA damage activation 24 h upon ATR inhibition, compared to resistant cells.

To determine whether the high expression of origin firing factors in sensitive cell lines led to a concomitant increase in replication origin firing, we analyzed the frequency of DNA replication origins per megabase of DNA (origin density) in our DNAscent data. There was no significant difference in origin density between sensitive and resistant cells in untreated conditions (**Figure 3g**). As expected, ATRi treatment for 1 hour resulted in increased origin firing in most of the cell lines, in line with the known role for ATR in suppressing origin firing. However, the increase in origin and fork densities upon 1 hour ATRi appeared overall higher in the sensitive cell lines and decreased after 24h, as compared to resistant cell lines (**Figure 3g-h, Extended Figure 3e**).

To test whether this hyper-increase in ATRi-induced origin firing was responsible for ATRi sensitivity, we sought to manipulate origin firing rates. We used XL-413, an inhibitor of CDC7, a positive regulator of origin firing^10^, henceforth termed ‘CDC7i’. In the absence of ATRi, CDC7i reduced origin firing, as measured by the DNAscent origin density and by imaging EdU incorporation, and increased fork speeds in sensitive HCC1806 cells, as expected (**Figure 3i-k**)^37^. CDC7i also rescued the ATRi-dependent effects on origin firing and replication fork speed in the third sensitive cell line HCC1806 (**Figure 3i-k**). Importantly, CDC7i also reduced the anti-proliferative effect of ATRi in both sensitive cell lines tested (**Figure 3l-m, Extended Figure 3f**), whereas in the two resistant cell lines tested there was either no effect, or a sensitization to ATRi (**Extended Figure 3g-h**). Overall, reducing origin firing capacity with CDC7i rescues slow fork speeds and ATRi sensitivity (**Figure 3i-k**).

We reasoned that the higher propensity to increase origin firing could render these cell lines more sensitive to ATRi through exhaustion of DNA replication factors, leading to replication catastrophe and cell death^38,39^. Accordingly, despite no significant change in the number of γH2AX foci in S-phase cells across the cell line panel upon ATRi (**Extended Figure 3i**), total γH2AX intensity was higher in two sensitive cell lines (HCC1806, HCC1569) (**Extended Figure 3j**). We also found a trend towards increased proportion of interphase cells exhibiting pan-nuclear γH2AX staining, a potential marker of replication catastrophe^39,40^ (**Figure 3n-p**) in the two sensitive cell lines tested, which could contribute to the observed increased cell death upon ATRi treatment (**Figure 3q**). BT-549 was the least sensitive of the three sensitive cell lines and exhibited a fork velocity similar to the resistant cell line panel (**Figure 1d, 2b-c**). Interestingly, BT-549 did not have a high-level of pan-nuclear γH2AX induced by ATRi, when compared to HCC1569 and HCC1806. This could suggest that BT-549 might not have reached the critical excess of origin firing leading to cell death. Overall, these data suggest that an increased capacity for origin firing in sensitive cell lines promotes lethal levels of origin firing and subsequent cell death upon ATRi treatment.

### A transcriptomic signature of DNA replication initiation correlates with sensitivity to ATRi across cell lines from different cancer types

We next explored whether we could use this novel mechanistic insight to predict ATRi sensitivity from gene expression information, a readily available data type that is also suited to clinical biomarker application. We first asked whether we could predict ATRi sensitivity based on increased origin firing capacity markers in a larger panel of breast cancer cell lines, taking advantage of publicly available breast cancer CCLE transcriptomic data with known IC50 values to ceralasertib (concentration of the inhibitor that inhibits cell survival by 50%). We performed a Gene Set Enrichment Analysis (GSEA) to compare ceralasertib-sensitive and resistant breast cell lines. Among the 200 most enriched pathways from gene ontology biological process terms, we counted the number of pathways related to each category and measured the average normalized enrichment score (NES). In sensitive cell lines, we found enrichment of mitosis related pathways as well as DNA replication (**Figure 4a**). Given our findings regarding origin firing capacity above, we also analyzed the enrichment of the ‘DNA replication initiation’ signature which was highly enriched in breast cancer (NES 1.74, p=0.003) (**Figure 4b**). This suggests that there is a gene expression footprint of the higher origin firing capacity in sensitive cell lines.

**Figure 4.**
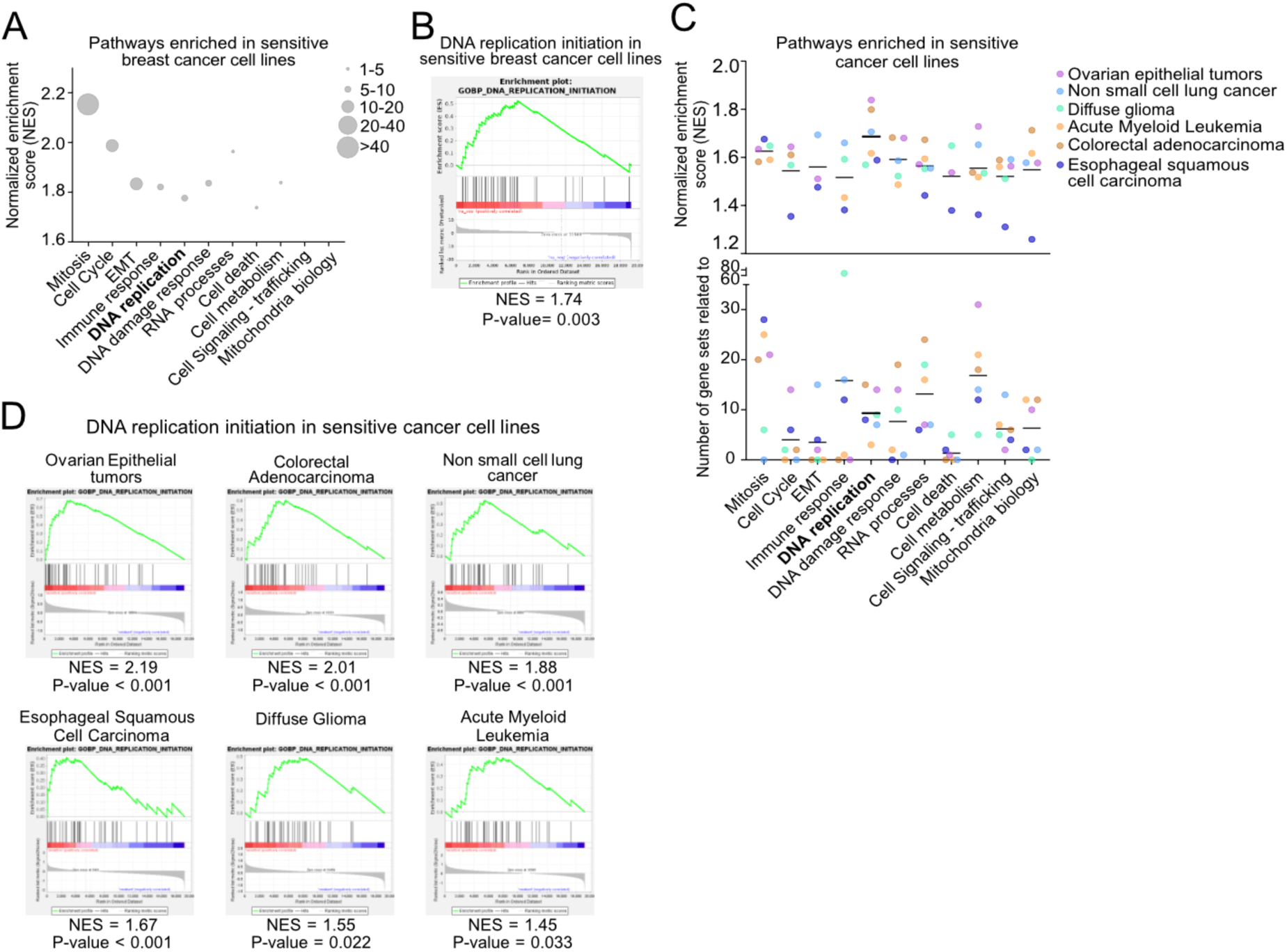
Transcriptomic signature of DNA replication initiation correlates with sensitivity to ATRi across cell lines from different cancer types. (**A**) Gene Set Enrichment Analysis (GSEA) analysis of publicly available transcriptomic data showing the normalized enrichment score (NES) and the number of pathways enriched per gene ontology (GO) biological processes in sensitive vs resistant breast cancer cell lines. (**B**) Enrichment of the DNA replication initiation signature from GO-terms biological processes in sensitive compared to resistant breast cancer cell lines. (**C**) GSEA of publicly available transcriptomic data showing the NES and the number of pathways enriched GO biological processes in sensitive vs resistant cancer cell lines. (**D**) Enrichment of the DNA replication initiation signature from GO-terms biological processes in sensitive cell lines compared to resistant cell lines. Sensitivity and resistance were determined using the top and bottom 25% of the ceralasertib IC50 from the GDSC2 dataset, normalized enrichment scores (NES) and nominal p-values were measured through the GSEA analysis.

Clinical trials involving ATRi are being carried out in a wide range of cancer types^11^. Therefore, we next sought to determine if gene expression signatures could also be indicative of ATRi sensitivity in additional cancer types beyond breast cancer. We performed a GSEA of sensitive and resistant cancer cell lines from CCLE from different cancer types, using publicly available RNA-sequencing data and IC50 data from the GDSC2 dataset. In this pan-cancer analysis, the highest NES across multiple cancers were for the DNA replication-related pathways (**Figure 4c**). Moreover, we found that the more specific ‘DNA replication initiation’ signature was significantly enriched in ceralasertib-sensitive cell lines from multiple cancer types with a particularly high enrichment in ovarian epithelial tumors and colorectal adenocarcinoma (**Figure 4d**). This reveals that origin firing factor gene expression may provide a new pan-cancer determinant of ATRi sensitivity that could be explored in patient samples.

### High origin firing capacity as a predictive marker of response to ATR inhibition in AML patient samples

Finally, we investigated whether our newly identified transcriptomic and proteomic determinants of ATRi sensitivity could be validated in cancer patient samples for potential clinical application. A previous study performed transcriptomic and proteomic analyses from AML patient samples. In addition, this study derived primary cell lines from patient samples, which were subject to a drug screen that included ceralasertib^35^. Using these published data, we identified 9 resistant and 8 sensitive primary cell lines, based on their measured IC50 (sensitivity below the first quartile; resistant above the third quartile) and investigated the related patient data for gene expression and kinase activities (**Figure 5a**). We first performed an unbiased GSEA (**Figure 5b**) in AML patients and found that even though mitosis related pathways were most represented (more than 40 pathways among 200), the DNA replication related pathways were the most enriched in sensitive samples (average NES = 2.2 vs NES = 2 for mitosis-related pathways). Additionally, we found that the ‘DNA replication initiation’ signature was even further enriched (p<0.0001, NES = 2.83) (**Figure 5b**). We next analyzed the phosphoproteomics data from the AML patient samples and discovered an enrichment of the kinase activities of CDC7 and CDK2 in ATRi sensitive samples (**Figure 5c**), exemplified by the increase of the phosphorylation sites MCM2-Serine 40 and Serine 139 involved in origin firing^41^ (**Figure 5d-e**). Altogether, our AML patient data analyses recapitulated ATRi sensitivity predictors discovered in cancer cell lines; the elevated gene expression of the ‘DNA replication initiation’ signature, and increased activity of kinases involved in origin firing, similar to the breast cancer phosphoproteomics analyses (**Figure 3**).

**Figure 5.**
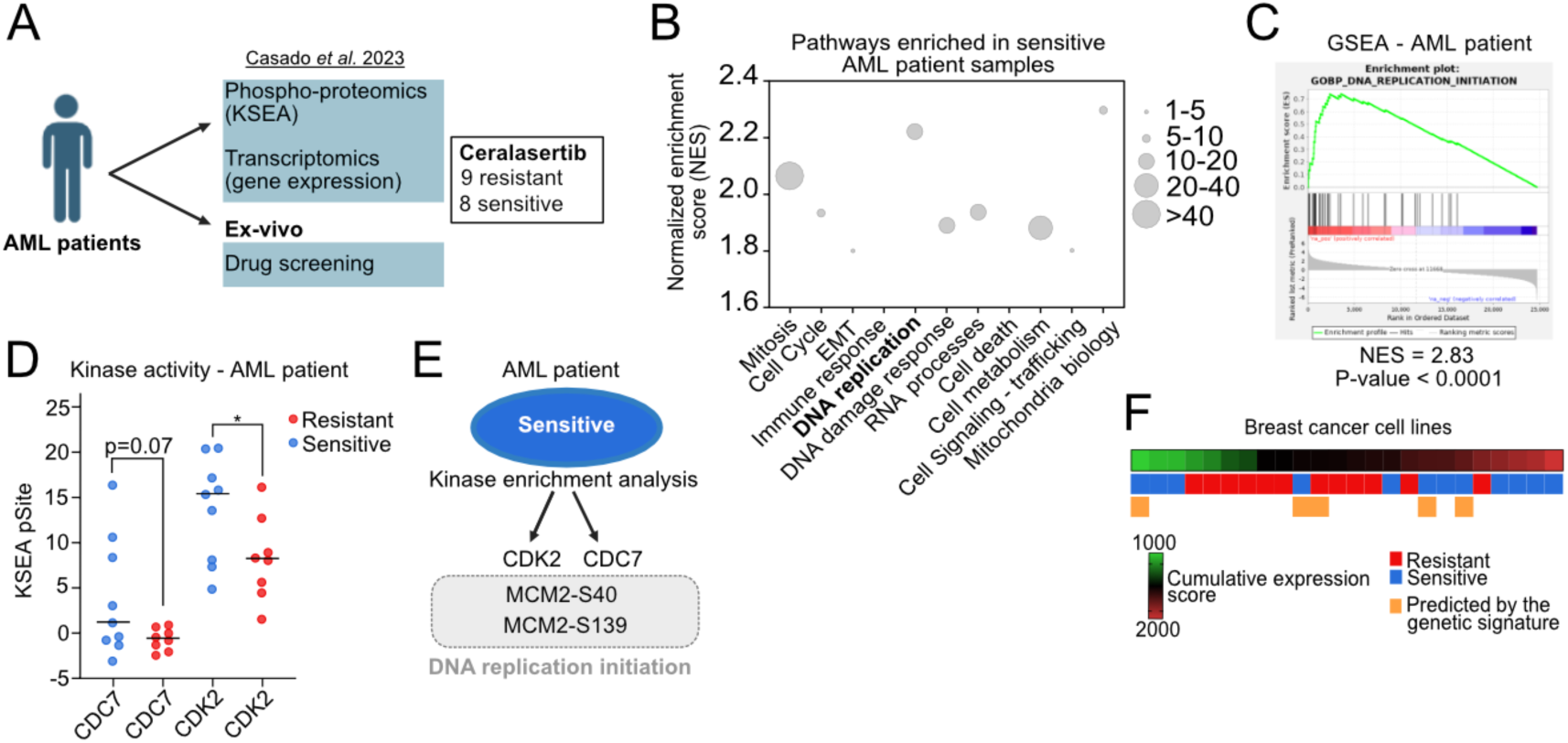
Exacerbated origin firing as a predictive marker of response to ATR inhibitor in cancer samples. (**A**) Schematic workflow of the data used from Casado et al. 2023 for analysis. Mass spectrometry and gene expression data are obtained from AML patient cells as described before^42^ and sensitivity to ceralasertib was obtained via ex-vivo drug screening. 9 samples had an IC50 higher than the third quartile of the dataset, qualified as resistant, and 8 samples lower than the first quartile, qualified as sensitive. **(B**) Gene Set Enrichment Analysis (GSEA) analysis of acute myeloid patient (AML) transcriptomic data showing the normalized enrichment score (NES) and the number of pathways enriched, per gene ontology (GO) biological processes, in predicted good responders compared to non-responders AML samples based on patient-derived primary cells sensitivity. (**C**) GSEA analysis of the gene ontology signature of DNA replication initiation in AML patient responders compared to non-responders to ceralasertib (NES = normalized enrichment score, nominal p-value). (**D**) Quantification of the Kinase Enrichment Analysis (KSEA) showing the enrichment of Kinase activities CDC7 and CDK2 in AML patients good responders compared to non-responders to ATRi. Unpaired t-test analysis. (**E**) Schematic showing the enrichment of CDK2 and CDC7 substrates at MCM2 sites S40 and S139 involved in origin firing. (**F**) Comparison of two predictive signatures; Heatmap of the cumulative expression score of the genes involved in the DNA replication initiation signature, in breast cancer cell lines, with track below indicating the sensitivity and the presence of cell lines in genetic cluster 3 predicted by copy number features of replication stress. *, P < 0.05.

In this study we identified two potential new determinants of sensitivity to ATR inhibition: a DNA CNA signature and a gene expression signature of DNA replication initiation. Interestingly, and perhaps reflecting the presence of multiple distinct pathways to ATRi sensitivity, we observed that there was not a clear overlap between the cell lines with the highest gene expression of origin firing factors and the ones predicted by the genetic signature of RS, with most predicted sensitive cell lines being signature mode-specific (**Figure 5f**). This suggests that our multi-omics study of the biology of ATRi revealed two likely different mechanisms of sensitivity to ATRi that have the potential to be further developed into a combined biomarker to select the patients most likely to benefit from ATRi therapies.

## Discussion

In this study, we set out to discover why some cancer cells are more sensitive to ATRi than others. Using a combination of genomic alteration patterns derived from prior work, and new functional genomics approaches herein, we identified two new determinants of sensitivity to ATR inhibition: exacerbated levels of origin firing, and a copy number signature of RS, that each identified largely non-overlapping sensitive cancer cell lines.

In normal immortalized cells (RPE1, FNE1), increasing RS rates with aphidicolin or cyclin E overexpression induced CIN and sensitivity to ATRi. However, cancer cells often have pre-existing CIN and complex genetic backgrounds, making it more difficult to predict sensitivity to ATR inhibition based on simplistic measures of RS or CIN. Indeed, ATRi treatment induced RS and CIN in all breast cancer cell lines tested regardless of sensitivity. Moreover, it was previously shown that nucleoside supplementation can dampen RS^32^ and CIN^28^, but this only mildly rescued ATRi sensitivity in our panel. Thus, we investigated additional features of RS to understand the differences in sensitivity between the breast cancer cell lines. CIN rates did not predict sensitivity *per se*, but the presence of specific CNAs close to large genes, an indicator of RS-induced CIN^17^, predicted a subset of sensitive cell lines. While this specific genomic signature is present in several sensitive cell lines, it must be further investigated in larger datasets as well as patient samples, as a potential biomarker of ATRi response.

Next, we found a correlation between slow replication fork velocity and sensitivity to ATR inhibition, though this was not enough to predict sensitivity to ATRi. Our data showed that ATRi sensitive cell lines had high expression and activation of origin firing factors, which correlated with hyper-increased origin firing in response to ATR inhibition. It is known that fork progression is slowed in the context of excessive origin firing^37^, potentially explaining this association. This excessive origin firing likely induces cell death in sensitive cell lines, while resistant cells restrain levels of origin firing at rates that do not affect their cell proliferation. Accordingly, reducing origin firing using CDC7 inhibition rescued ATRi-induced alterations in origin firing and replication fork speed. CDC7 inhibition also partially rescued ATRi proliferation defects in sensitive cell lines.

Our data suggest replication catastrophe as the link between excessive origin firing, pan-nuclear gH2AX staining, and cell death. It is important however to consider the difficulty of robustly detecting cell death by replication catastrophe; while we use pan-nuclear γH2AX staining as an indicator, the exact mechanisms are not well known, making an unambiguous marker elusive. RPA staining could also be used to understand whether replication catastrophe is caused by RPA exhaustion because of excessive origin firing in our cell lines^38,39^, as nucleotides do not seem to be a limiting factor. In normal immortalized cell lines, we found that cyclin E overexpression sensitized cells to ATRi. This did not seem to be the general mechanism of sensitivity in breast cancer cell lines, although *CCNE1* amplification has been described in the sensitive line HCC1806^43^. A recent study suggested that ATRi sensitivity correlated with the induction of γH2AX signals and hyper-activity of CDK2 via unknown mechanisms^44^. Our study now places these findings in the context of an upregulated origin firing pathway being the key determinant of ATRi sensitivity.

We sought to determine if we could further apply this mechanism of ATRi action as a potential biomarker to predict ATRi sensitivity across additional breast cancer cell lines, and across multiple cancer types. We performed transcriptomic analysis and found recurrent pathways enriched in sensitive cancer cell lines compared to resistant related to mitosis, DNA replication, immune response and cell metabolism. Based on the data gathered in this study, we closely examined the DNA replication pathways and found that ATRi sensitivity in CCLE cell lines from different cancer types and AML patients mirrored what we observed in our breast cancer cell line panel: high expression of origin firing factors correlated with increased ATRi sensitivity. Interestingly, we also found evidence of higher origin firing kinase activities (CDC7, CDK2 specific activities) in AML patient samples sensitive to ATRi. Investigating the combination of the transcriptomic signature with a measure of activity of origin firing kinases could be used to develop a biomarker tool and improve patient selection strategies. Although we focused in this study on the RS-related pathways revealed by our analyses, we also noted that mitosis related pathways were highly enriched in sensitive breast and esophageal squamous cancer cell lines, as well as mildly enriched in the other cancer types and in AML patient samples. The relationship between CDK1 activity, mitotic processes and sensitivity to ATR inhibition is not the subject of this study but could lead to further investigation as it could help increase the understanding of ATRi biology and unravel new determinants of sensitivity.

The mechanistic insight of our study provides a first step in developing multimodal biomarkers for ATRi and provides information to assist in designing combination treatment strategies. For example, it may be possible to sensitize ATRi resistant patients by manipulating origin firing capacity. Interestingly, it was previously shown that CDC7i and ATRi combination have a synergistic effect in hepatocellular carcinoma^45^, by decreasing cancer cell viability and mouse tumor growth, suggesting this combination might be useful to sensitize resistant tumors. Of note, we found that one resistant cell line indeed was sensitized to ATRi upon CDC7 inhibition. This suggests that sensitive cell lines with high origin firing capacity could benefit from ATRi monotherapy, while resistant tumors may be targeted using a combination of ATRi and CDC7i.

Finally, having identified two independent indicators of ATRi sensitivity, we wondered if these would select the same, or different subsets of ATRi sensitive cell lines. We observed that ATRi sensitive cell lines identified by the genomic signature were generally exclusive with the ATRi sensitive cell lines exhibiting the highest expression of the DNA replication initiation signature. This indicated there may be (at least) two distinct mechanisms predictive of response to ATR inhibition, and that by combining these transcriptomic, copy number and phosphoproteomics signatures, a multi-modal biomarker could be developed. Indeed, such a combinatory approach would increase the number of sensitive tumor samples detected but also decrease the number of false positives. In this study we were limited to the few available datasets that included RNA-sequencing, genomics, mass spectrometry and sensitivity (IC50) data. Therefore, future investigations on larger scale multi-omics datasets linked to ATRi efficacy are needed to translate this knowledge into a potential biomarker suitable for clinical applications.

## Material and methods

### Tissue culture

All cell lines were grown in their recommended media supplemented with 1X Anti-Anti (Gibco™, 15240-062) and 10% heat inactivated Fetal Bovine Serum (Gibco™, 10082147) and were incubated at 37 degrees with 5% CO2 supply. HCC1569, HCC1806 and HCC1187 are cultured in RPMI 1640 Medium (Gibco™, 21875-034); T47D and BT-549 in RPMI supplemented with 8.3344 µg/mL and 0.958 µg/mL of Insulin-Transferrin-Selenium (ITS-G) (Gibco™, 41400-045) respectively; MDA-MB-231 in Dulbecco’s Modified Eagle Medium (Gibco™, 41966-029); BT-474 in Hybri-Care Medium (ATCC™, 46-X.) supplemented with 1.5 g/L sodium bicarbonate; and BT-20 in Minimum Essential Media (Gibco™, 21575-022). Cell lines were obtained from AstraZeneca and the Francis Crick Institute, STR-profiled and Mycoplasma tested.

FNE1 immortalized (hTERT) fallopian tube cells were obtained from University of Miami through an MTA and cultured in FOMI media^20^. RPE1 cells were purchased from ATCC® and grown in F-12 Nutrient Mix (Ham, 1X) (Gibco™, No. 21765-029). KTB6, KTB21, KTB34, and KTB6 immortalized (hTERT) normal breast cells were cultured in 25% Gibco™ low glucose DMEM (Gibco™, No. 12320-032) with 75% F-12 Nutrient Mix (Ham, 1X) (Gibco™, No. 21765-029) supplemented with 0.4 µg/mL hydrocortisone (Sigma-Aldrich, H0888), 5 µg/mL of Insulin (Sigma Aldrich, I5500), 20 ng/mL EGF (Sigma Aldrich, E9644), 0.5 µl/ml of Y-27632 (Rock inhibitor, ALX-279-333-M005, Enzo Life Sciences) and 4 µl/mL of Adenosine (Sigma-Aldrich, A-9001)^24^. Cells were obtained from The Susan G. Komen Tissue Bank (KTB) at Indiana University Simon Comprehensive Cancer Centre (IUSCCC) through an MTA (material transfer agreement).

### Cell Confluence Assay

Cells were seeded in 96-well plates at a range of 3,000-10,000 cells per well depending on the cell line. 24 hours after seeding, the medium was changed, and cells were treated with a dose range of ceralasertib 0.05 µM - 3.2 µM. Phase contrast images were acquired every three hours for at least 72 hours using an Incucyte® S3 (Sartorius). Incucyte® software was used to apply a cellular mask for each cell line measuring the percentage cell confluence. Each experiment was performed in at least three independent replicates.

### Immunofluorescence

Cells were grown on glass coverslips in 6-well plates and fixed with PTEMF as described before^17^. After blocking with 3% BSA, cells were stained with primary antibodies against p-H2AX S139 (Millipore 05- 636, 1:500) and CREST (antibodies incorporated, 15-234-0001, 1:500), and with secondary antibodies goat anti-mouse Alexa Fluor 488 (A11017 Invitrogen, 1:500) and goat anti-human Alexa Fluor 647 (A21445 Invitrogen, 1:500). DNA was stained with DAPI (Roche) and coverslips were mounted using Vectashield (Vector H-1000, Vector Laboratories). For EdU (5-ethynyl-20-deoxyuridine) experiments, cells were incubated for 15 minutes with 10 µM EdU (Thermo Fisher Scientific) prior to fixation. After fixation, EdU was stained using Click-iT EdU imaging kit (Invitrogen, C10338) as indicated in the manufacturer’s instructions.

Images were acquired using the Olympus DeltaVision RT microscope and deconvolved with SoftWorx Explorer as before^17^. For segregation errors, cells were imaged at 100x objective; for micronuclei quantification at 40x objective; and at 20x for cell cycle analysis and γH2AX signals. Analysis of immunofluorescence images was performed using ImageJ and CellProfiler software.

### Genomic clustering – CNA analysis

Breast cancer cell line whole genome sequencing data was downloaded from CCLE. A panel of normal was collated from the 1000 human genome project, using only female samples, to control for artefacts. For both sample sets QC and adaptor trimming was performed using Trimgalore and aligned to GRCh38 using bwa mem. Then duplicate reads were removed using Picard. To estimate somatic copy number alterations CNVkit was implemented with default parameters. For all CNAs the interstitial breakpoint positions the distance to the nearest large gene was calculated. A list of large genes, gene cut-off of 600kb, was assembled from UCSC genome browser – GRCh38. Each CNV feature (loss, gain and distance to large genes) was log normalised if required, and scaled from 0-1, before performing hierarchical clustering using Euclidean distances and Ward’s linkage method for all samples. We cut the dendrogram into three clusters. Then the IC50 of each cluster was interrogated from Genomics of drug sensitivity and Cancer (GDSC2) database^27^.

### Replication fork dynamics from ONT sequencing and DNAscent

Cells were incubated with 50 µM EdU for 5 minutes, 50 µM BrdU for 10 minutes, followed by thymidine for 20 minutes, with washes performed between each incubation. Cells were collected by scraping into cold PBS, pelleted and washed at 4 degrees and frozen at -80 degrees. High molecular weight genomic DNA was extracted using the QIAGEN® Puregene Core kit B (No. 1042608). The concentration, quality and length of DNA were measured using Tape Station and sample libraries were prepared using Nanopore kits SQK-LSK109 for R9.4.1 and SQK-ULK-114 for R10.4.1, quality controlled via Tape Station and sequenced with the ONT PromethION platform. Library preparation and sequencing was performed by the genomics facility of the Universities of Birmingham and Cambridge.

For samples sequenced on the ONT PromethION platform with a R9.4.1 nanopore (TBD kit details for all samples) base calling of raw sequencing files (fast5 format) was performed using GUPPY (version 5.0.16) with the respective configuration (dna_r9.4.1_450bps_hac_prom.cfg). Both reads that passed and failed the base calling quality metrics were included in downstream processing. Reads were aligned to the human telomere-to-telomere (T2T) reference genome (chm13v2.0.fa) using minimap 2 (version 2.24). Next, the output was converted from sam format to bam format using samtools (version 1.14). Finally, the DNAscent pipeline (version 3.1.2) with subprogrammes index, detect, forkSense was run as previously described^31^. The DNAscent forkSense output (bed file format) was processed with custom Python (version 3.11.5) scripts to obtain fork speeds and stall scores. During this study the R9.4.1 flow cells were discontinued so experiments with HCC1806, Cdc7i and ATRi were performed on the ONT PromethION platform with a R10.4.1 nanopore. Base calling of raw sequencing files (pod5 format) and alignment (--mm2-preset map-ont option) were performed using Dorado (version 0.7.3) and the configuration dna_r10.4.1_e8.2_400bps_fast@v5.0.0 and the T2T reference genome (chm13v2.0.fa).

The DNAscent pipeline (version 4.0.3, commit 1432cf75f1e57920856f1ca465ad3bc5650ebbe9, available on https://github.com/MBoemo/DNAscent) with subprogrammes index, detect and forkSense was run and the output was processed in the same way as described above for the R9.4.1 pore (DNAscent v3.1.2). To generate origin and fork density metrics, the total numbers of origins and forks in the sample were divided by the length of megabase sequenced to obtain a density per megabase.

### Copy number profile

Pod5 raw sequencing files were basecalled using Dorado v0.5.3. Adaptors were trimmed from fastq files using Porechop v0.2.4 and aligned to GRCh37 using minimap2 v2.5. Segmented copy number data was generated using 500kb bins using QDNASeq followed by read count, mappability and GC content correction and then copy number segmentation. Absolute copy number fitting was performed using ACE using cellularity, ploidy and error estimations from the squaremodel function.

### Sample preparation and TMT labelling

Cells were washed with PBS and scraped in SDS 4%, 50 mM Tris HCl pH 7.4 for cell lysis and protein extraction. After sonication and lysate clearing by centrifugation, protein concentration was measured using BCA protein assay kit and balanced to 55 µg for every sample.

To prepare the samples for proteomics analysis, isobaric filter-aided sample preparation was performed^46^. Briefly, samples were diluted in UA buffer (8 M Urea, 100 mM Tris-HCl pH 8.5 in dH2O) and reduced with the addition of 10 mM DTT (VWR, M109) and incubated for 30 min at room temperature. Samples were then alkylated with 55 mM IAA (Iodoacetamide, VWR, 786–228) and incubated in the dark for 30 min at room temperature. Samples were loaded on Vivacon® 500, 30,000 MWCO Hydrosart filters (Sartorius, VN01H22) and centrifuged at 14,000 RCF for 20 min. Sampled were washed twice with UA buffer and centrifuged at 14,000 RCF for 20 min. Then, samples were washed three times with 100 mM TEAB (Triethylammonium bicarbonate, Fisher Scientific, 15215753) before a final addition of 100 μL 100 nM TEAB and 1 μg of Trypsin (Merck, T6567) to every sample before an overnight incubation at 37C.

Each TMTpro™ 18-plex Label Reagent (ThermoFisher Scientific, A52047) was resuspended in 100% Acetonitrile and 8μg of TMT-Pro label was added to each respective sample filter before incubation for 1 hour at 25°C at 600 RPM. Excess TMT-Pro labels were quenched with the addition of 5% hydroxylamine and incubated for another 30 mins. Peptides were eluted by centrifugation at 14,000 RCF for 20 mins, the filters were washed using 100 mM TEAB and a final wash of 30% acetonitrile. All labelled peptides were pooled and split 10% for the total proteomics analysis and 90% for the phospho-enrichment. The pooled peptide mixture was dried with a vacuum concentrator before downstream sample preparation.

For total proteomics analysis, the TMT-labelled pool was fractionated into seven fractions using Pierce™ High pH reverse-phase fractionation kit (Life Technologies, 84868), using the manufacturers protocol. Fractions were dried using vacuum concentration before being resuspended in Buffer A* (2% ACN, 0.1% TFA, 0.5% Acetic Acid) for LC-MS/MS analysis. For phosphoproteomics analysis, phosphopeptides were enriched for using Titansphere TiO phospho-peptide enrichment kit (GL Sciences, 5010-21308), following the manufacturer’s instructions. Eluted enriched TMT-labelled phosphopeptides were dried using vacuum centrifugation before resuspension in Buffer A* for LC- MS/MS analysis.

### Mass Spectrometry analysis and data processing

LC–MS/MS analysis was performed on a Q Exactive-plus Orbitrap mass spectrometer coupled with a nanoflow ultimate 3000 RSL nano HPLC platform (Thermo Fisher). Dried peptide mixtures were resuspended in A* buffer. For total proteomics analysis, equivalent of ∼1 μg of protein was injected into the nanoflow HPLC. For phospho-proteomics analysis, ∼90% of the total peptide mixture was injected. Samples were resolved at flow rate of 250 nL/min on an Easy-Spray 50 cm × 75 μm RSLC C18 column (Thermo Fisher). Each run consisted of a 123 mins gradient of 3% to 35% of Buffer B (0.1% FA in Acetonitrile) against Buffer A (0.1% FA in LC–MS gradient water), and separated samples were infused into the MS by electrospray ionization (ESI). Spray voltage was set at 1.95 kV, and capillary temperature was set to 255°C. MS was operated in data-dependent positive mode, where 1 MS scan is followed by 15 MS2 scans (top 15 methods). Full-scan survey spectra (m/z 375–1,500) were acquired with a 70,000 resolution for MS scans and 35,000 for the MS2 scans. A 30-s dynamic exclusion was applied.

Mass spectrometry raw data files were searched and quantified against a FASTA file of the Homo sapiens proteome using MaxQuant (version 2.2.2.0)^47^. Parameter groups were set to define the phospho- or total-proteome analysis and fractions were set for the total proteomics analysis. “Reporter ion MS2” type option was selected with a reporter mass tolerance of 0.003 Da. Enzyme specificity was set to “Trypsin,” allowing up to two missed cleavages. Phosphorylation (STY) was added to variable modifications for the phospho-parameter group. False discovery rates (FDR) were calculated using a reverse database search approach and were set at 1%. “Re-quantify” options were enabled. Default MaxQuant settings were used for all other parameters.

Downstream statistical analysis was performed in Perseus (version 1.6.2.3)^47^ where data was filtered for contaminants, reverse peptides and peptides only identified by site, log2 transformed, and filtered for valid values only. The Log2 data generated were used for subsequent analysis of total protein expression and Kinase Enrichment Analysis (KSEA).

### KSEA analysis

Phosphoproteomics Log2 transformed fold changes were translated into kinase activity using Kinase Enrichment Analysis (KSEA) as previously described^35,48^, available on https://github.com/CutillasLab/protools2 using signor and Phosphositepus as input databases of kinase-substrate relationships^49,50^. The KSEA analysis was performed to compare a) three resistant DMSO and three sensitive DMSO to reveal basal differences of kinase activities and b) between ATRi treated samples (1h and 24h) and their relative DMSO to reveal kinase activities induced by the treatment. Changes in kinase activity were inferred from variations in the putative downstream targets (pdts) of each kinase the significance of these changes were calculated using Kolmogorov–Smirnov tests^51^. Z-score were plotted to visualize the most enriched kinase activities between the different conditions. Volcano plots were generated using the volcano plot function on R using z-scores and p-values.

### Transcriptomic analysis

Publicly available RNA-sequencing data of the Cancer Cell Line Encyclopedia (CCLE, broadinstitute.org) were downloaded as transcript per million. IC50 of the cancer cell lines were obtained from the GDSC2 dataset and used to classify the cell lines as most sensitive or most resistant (IC50 lower than the first quartile or higher than the third quartile respectively). Gene set enrichment analysis (GSEA) was performed between the sensitive and the resistant cell line groups using Gene Ontology – Biological Processes (GO-BP) (c5.go.bp.v2024.1) gene sets as reference in the GSEA software (Broad Institute). Normalized Enrichment Score (NES) and nominal q-values were used to identify the significantly enriched GO-term BP in sensitive cell lines.

The transcriptomic data of a cohort of AML patients, classified as good responders or not responders to ceralasertib, were obtained from Casado and colleagues study^42^. Genes were ranked in descending order of fold change (FC) and a gene set enrichment analysis was performed using the ranking module, between the good responders and non-responders to the ATR inhibitor. Gene Ontology – Biological Processes (GO-BP) (c5.go.bp.v2024.1) gene sets were used as reference.

### Statistical analysis

Statistical analyses were performed using GraphPad Prism 10.1.2 (324). To compare two datasets, we performed normality tests using Agostino and Pearson and Shapiro-Wilt tests. Paired or unpaired t-test, Mann and Whitney and Wilcoxon rank tests were performed depending on the data distribution and type. For more than two datasets, the One-way ANOVA test was done. In all cases, ns indicates not significant; *, P<0.05; **, P<0.01; ***, P<0.001; ****, P<0.0001. More information can be found in the Figure legends.

## Data availability

Gene expression, long-read sequencing and mass spectrometry analysis raw data generated in this study can be found in the file ‘Supplementary Tables’ Excel spreadsheet. The long-read sequencing data for this study were deposited in the European Nucleotide Archive (ENA) at EMBL-EBI under accession number PRJEB85428. The mass spectrometry raw data files were deposited on ProteomeXchange Consortium via the PRIDE partner repository.

## Acknowledgements

The authors would like to thank AstraZeneca for providing breast cancer cell lines and funding this project. We would like to thank Alan Lau for his input on the manuscript. Next, we would like to thank the Cambridge Genomics Services at the Department of Pathology and the Birmingham Genomics Services at the Institute of Cancer and Genomic Sciences for the Oxford Nanopore Technologies long-read sequencing. This work was performed using resources provided by the Cambridge Service for Data Driven Discovery (CSD3) operated by the University of Cambridge Research Computing Service (www.csd3.cam.ac.uk), provided by Dell EMC and Intel using Tier-2 funding from the Engineering and Physical Sciences Research Council (capital grant EP/T022159/1), and DiRAC funding from the Science and Technology Facilities Council (www.dirac.ac.uk). We thank all the members of the McClelland lab for critical reading of the manuscript and general discussions. We are thankful to the Barts Cancer Institute microscopy and mass spectrometry facilities, for the support in the generation of microscopy images (Incucyte, DeltaVision) and mass spectrometry data. We would like to thank the Komen Tissue Bank (KTB) for sharing the KTB cell lines used in this paper. Several schematic images were made using the Biorender designing online tool.

## Authors contributions

AL designed the study, performed experiments, analyzed data and wrote the article, supervised by SEM. PLP performed DNAscent analyses and analyzed data, supervised by MB. JAS performed experiments related to Figure 2, supervised by SEM. SDA performed computational analyses relating to the copy number signature, supervised by SEM. FBC analyzed gene expression data and provided RNA-Seq data of AML patients. MAG analyzed data, supervised by SEM. GH analyzed data, supervised by PC. NS performed experiments relating to Figure 1b and provided advice, supervised by SEM. ELA performed mass spectrometry experiments, supervised by FKM. PRC analyzed the phosphoproteomics data and provided mass spectrometry data of AML patients. FKM analyzed data. JVF provided advice and data interpretation. MAB provided advice and data interpretation. SEM designed the study, supervised experiments and wrote the manuscript with input from all authors.

## Extended Figures

**Extended Figure 1.**
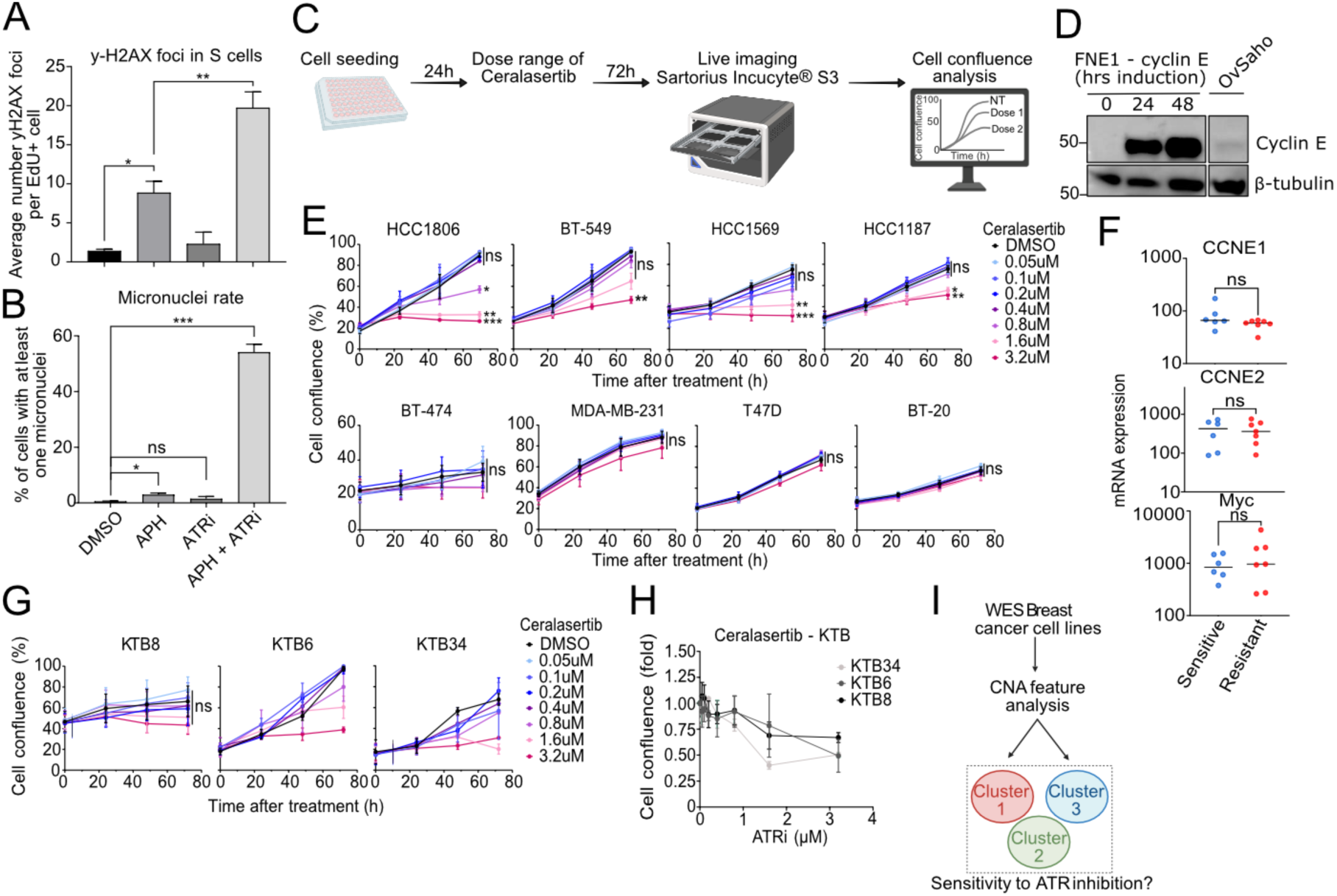
(**A**) Quantification of the average number of γH2AX foci in replicating cells (EdU+) and (**B**) of the micronuclei rate in RPE hTERT cells treated with DMSO, APH 0.4uM, ATRi 0.5 µM or both for 24h. Results were obtained from three independent immunofluorescence replicates. Paired t-test analysis. (**C**) Western Blot of Cyclin E and β-tubulin proteins in FNE1 cells overexpressing Cyclin E, induced with 2 µg/mL of doxycycline for 0h, 24h or 48h, and Ovsaho ovarian cancer cell line. (**D**) Representation of the protocol used to perform the sensitivity assay to ATR inhibitor using the incucyte microscopy. Cells were treated 24h after seeding and the cell confluence was measured every few hours for 72h. (**E**) Quantification of the confluence over time in response to different doses of ATRi in eight different breast cancer cell lines. Multiple paired t-test analysis. (**F**) Quantification of the mRNA expression (normalized transcripts per million) of *CCNE1, CCNE2* and *Myc* in sensitive and resistant breast cancer cell lines using publicly available data from CCLE. Sensitivity was determined using the GDSC2 dataset, the top 25% of cell lines were considered resistant and the bottom 25% as sensitive. Unpaired t-test analysis. (**G**) Cell confluence dose response of ceralasertib (ATRi) was measured at 72h post-treatment in three KTB immortalized cell lines. Fold confluence of ATRi treated samples was measured related to DMSO at 72h endpoint. Results are representative of at least two independent replicates. (**H**) Quantification of the confluence over time in response to different doses of ATRi in three different KTB breast cell lines. (**I**) Schematic representation of the DNA copy number clustering strategy based on Whole Exome Sequencing data of a large panel of breast cancer cell lines. Multiple paired t-test analysis. Ns, not significant; *, P < 0.05; **, P < 0.01; ***, P < 0.001.

**Extended Figure 2.**
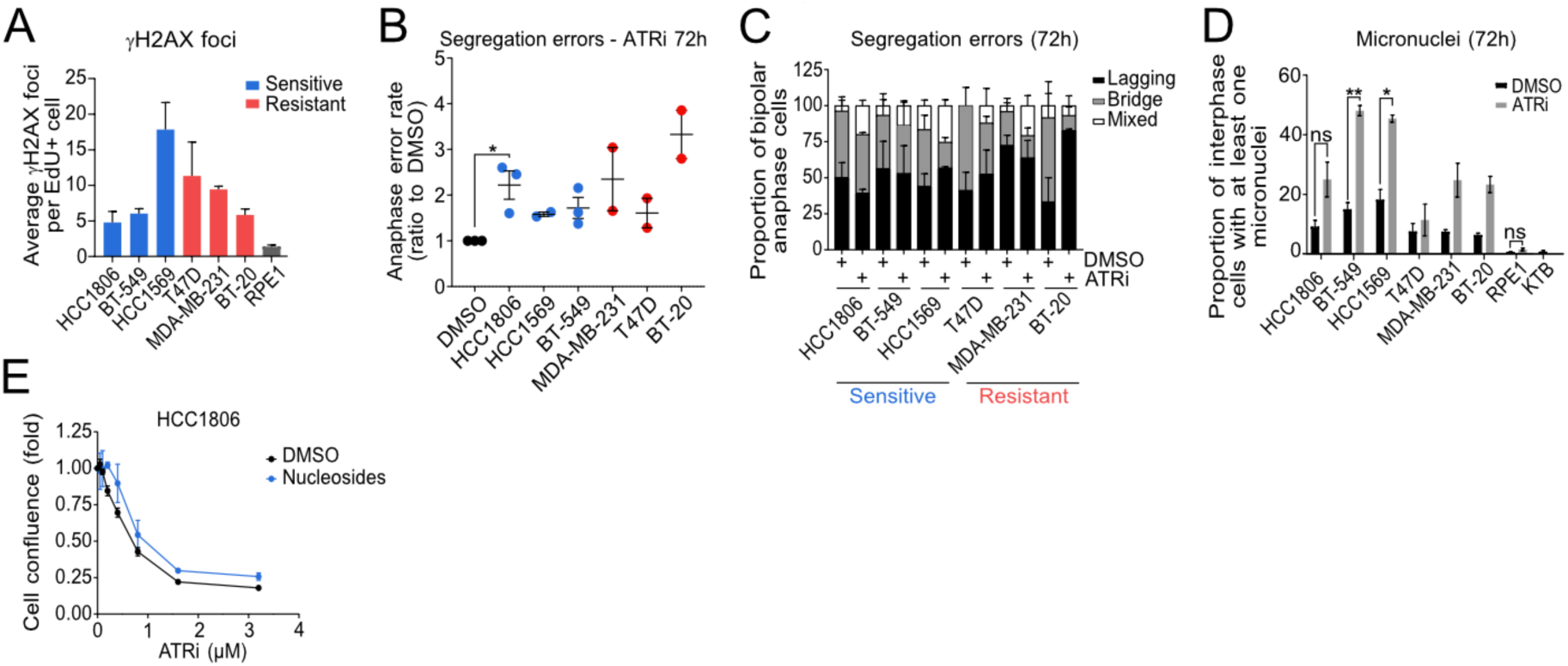
(**A**) Quantification of the mean number of γH2AX foci per replicating cell (EdU+) in breast cancer cell lines and RPE hTERT. Results are representative of at least two independent replicates per cell line. (**B**) Quantification of the segregation error rate of breast cancer cell lines in response to 72 hours of 0.5 µM ATRi, fold to DMSO. (**C**) Quantification of the segregation error types per anaphase, with the proportion of lagging chromosomes, chromosome bridges and mixed (lagging and bridge). (**D**) Quantification of the micronuclei rate in breast cancer cell lines treated with 0.5 µM ATRi or DMSO for 72 hours. Result of segregation error and micronuclei rates are representative of at least two independent replicates per cell line. Paired t-test analysis (**E**) Cell confluence dose response of ceralasertib (ATRi) in DMSO or 10X EmbryoMax® Nucleosides HCC1806 treated cells. Fold confluence of ATRi treated samples was measured related to DMSO at 72 hours endpoint. Results are representative of two independent replicates. Paired t-test analysis. Ns, not significant; *, P < 0.05; **, P < 0.01.

**Extended Figure 3.**
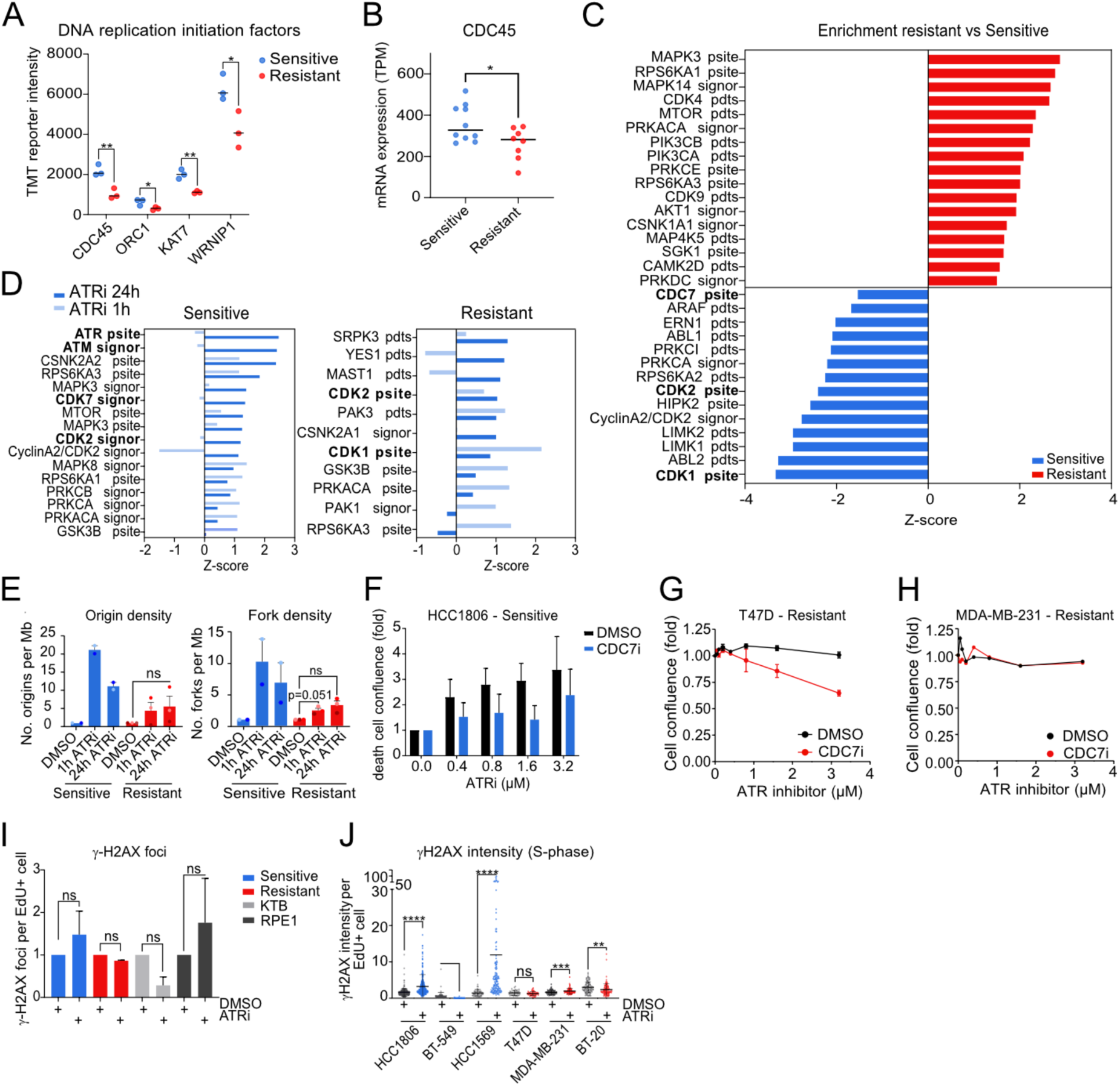
(**A**) TMT reporter (A.U.) of DNA replication initiation factors CDC45, ORC1, KAT7 and WRNIP1 in sensitive and resistant breast cancer cell lines. Unpaired t-test analysis. (**B**) Quantification of the mRNA expression of CDC45 gene in sensitive and resistant breast cancer cell line using publicly available data from CCLE. Unpaired t-test analysis. (**C**) Quantification of the enriched kinase activities (z-score) in Resistant compared to Sensitive breast cancer cell lines. In red are the kinase activities enriched in resistant and in blue in sensitive cell lines. (**D**) Quantification of the kinase activities enriched in sensitive and resistant cell lines enriched in 0.5 µM ATRi for 1 hour or 24 hours compared to DMSO. (**E**) Quantification of the replication origins and forks density per megabase, fold increase to DMSO, in breast cancer cell lines treated with DMSO, 0.5 µM ATRi for 1 hour or 24 hours. Data obtained from R9 long-read DNA sequencing. Paired t-test analysis. (**F**) Quantification of the proportion of death cell confluence using a green cytotox live death marker in HCC1806 cells in response to ATRi only or to ATRi and 0.25 µM CDC7i. Results are measured after 72 hours of treatment and the fold change is measured for each ATRi treatment to the DMSO and are representative of two independent experiments. (**G,H**) Quantification of the dose response to ATRi in combination with DMSO or 0.25µM of CDC7i in T47D (F) and MDA-MB-231 (G) resistant breast cancer cell lines. (**I**) Quantification of the number of γH2AX foci per replicating cells (EdU+) in breast cancer, KTB and RPE1 cell lines treated with DMSO or 0.5 µM ATRi for 24 hours. Paired t-test analysis. (**J**) Quantification of the γH2AX intensity per replicating cell (EdU+) in breast cancer cell lines treated with DMSO or 0.5 µM ATRi for 24 hours. Paired t-test analysis. Ns, not significant; *, P < 0.05; **, P < 0.01; ****, P < 0.0001.

